# Therapeutic Potential of Cyclodextrins Targeting Dengue Virus and SARS-CoV-2 Infection and Pathogenesis

**DOI:** 10.1101/2025.08.03.668365

**Authors:** Pedro H. Carneiro, E. Vanessa Jimenez-Posada, Francielle Tramontini Gomes de Sousa, Scott B. Biering, Trishna S. Patel, Samhita Bhat, Sarah Stanley, John Pak, Peter Laing, Tamas Sohajda, Eva Harris, P. Robert Beatty

**Affiliations:** Division of Infectious Diseases and Vaccinology, School of Public Health, University of California, Berkeley, Berkeley, CA, USA; Division of Immunology and Molecular Medicine, Department of Molecular and Cell Biology, University of California, Berkeley, Berkeley, CA, USA; Chan Zuckerberg Biohub, San Francisco, CA, USA; Excivion Ltd., Cambridge, United Kingdom; CarboHyde, Budapest, Hungary

## Abstract

Cyclodextrins (CDs) are cyclic oligosaccharides with promising therapeutic applications, including antiviral activity. During viral infections, pathogenesis arises not only from viral replication but also from viral proteins that act as “toxins”, disrupting cellular barriers and inducing endothelial dysfunction, a hallmark of severe diseases such as dengue and COVID-19. Dengue virus (DENV) NS1 and SARS-CoV-2 Spike proteins induce endothelial hyperpermeability, contributing to severe complications. Here we explored the potential of a panel of 18 CDs in mitigating endothelial dysfunction caused by these viral proteins and evaluated the CDs’ antiviral activity *in vitro* and *in vivo*. The effect of CDs on endothelial hyperpermeability was assessed using a trans-endothelial electrical resistance assay with human pulmonary microvascular endothelial cells exposed to DENV NS1 and SARS-CoV-2 Spike proteins. Antiviral efficacy of CDs was evaluated in Vero cells infected with DENV2 and Calu-3 cells infected with SARS-CoV-2, and in vivo protection was assessed in a lethal DENV2 mouse model. CDs effectively inhibited DENV NS1-induced endothelial hyperpermeability *in vitro*, demonstrating their potential to counteract NS1-mediated barrier disruption. In the murine model, CD1 treatment provided partial protection against DENV-induced morbidity and mortality. Further, CDs significantly reduced SARS-CoV-2 infection *in vitro* and inhibited Spike-induced endothelial dysfunction. These findings indicate that CDs can prevent endothelial hyperpermeability induced by DENV NS1 and SARS-CoV-2 Spike proteins and exhibit antiviral activity against SARS-CoV-2, positioning them as promising candidates for mitigating endothelial complications associated with viral infections. Further research is needed to explore the clinical relevance of CDs and their mechanisms of action.

## 1. Introduction

Cyclodextrins (CDs) are a family of cyclic oligosaccharides with an array of various chemical and clinical functions that are used as drug excipients (Jambhekar and Breen, 2016). CDs are characterized by their unique ability to form inclusion complexes with a variety of molecules, ranging from small organic compounds to large biomolecules, inside a hydrophobic central cavity with a hydrophilic outer surface (Di Cagno, 2016; Jambhekar and Breen, 2016). CDs are currently approved for clinical use as excipients, but also with potential use as active pharmaceutical ingredients (Di Cagno, 2016).

CDs have shown promise in combating viruses through various mechanisms (Braga et al., 2021; Di Cagno, 2016; Jambhekar and Breen, 2016; Jones et al., 2020). Sulfated CD derivatives, such as sulfobutylether-β-cyclodextrin (SBE-β-CD), have been reported to inhibit SARS-CoV-2 viral entry by blocking viral attachment or fusion with host cells (Alboni et al., 2023). Previous studies have demonstrated the ability of CDs to prevent the entry of enveloped viruses like HIV (Garrido et al., 2020; Matassoli et al., 2018) and SARS-CoV-2 (Bezerra et al., 2022) by interfering with the interactions of viral glycoproteins with cellular receptors.

Diseases such as dengue and COVID-19 exemplify the devastating impact of viral epidemics on communities globally, resulting in widespread morbidity, mortality, and socioeconomic disruption (Hussein et al., 2024; Katzelnick et al., 2017). Dengue virus (DENV), transmitted primarily by *Aedes* mosquitoes, is widespread in tropical and subtropical regions, causing recurrent epidemics of disease characterized by high fever, severe joint and muscle pain, and, in severe cases, hemorrhagic manifestations and vascular leak that can lead to death (Katzelnick et al., 2024). The spread of SARS-CoV-2 precipitated a global pandemic of COVID-19 unprecedented in scale and severity (Hussein et al., 2024). COVID-19 presents with a wide spectrum of symptoms, ranging from mild respiratory illness to severe pneumonia and acute respiratory distress syndrome, overwhelming healthcare systems and causing immense societal disruption (Hussein et al., 2024). The major global burden of both dengue and COVID-19 highlight the urgent need for therapeutics to mitigate the impact of viral infections.

Viral proteins play an important role in the pathogenesis of viral infections, exerting diverse effects on host cells and tissues, including the induction of vascular leakage. One such example is the DENV nonstructural protein 1 (NS1), which acts independently of the virus as a virulence factor, or viral “toxin”, contributing to the severity of dengue disease by disrupting endothelial barrier function, promoting vascular permeability, and exacerbating plasma leakage syndrome (Beatty et al., 2015; Modhiran et al., 2015; Puerta-Guardo et al., 2019, 2016). Similarly, besides its well-established role in viral entry into host cells, the Spike protein of SARS-CoV-2 has been shown to be a key virulence determinant, triggering endothelial permeability and promoting vascular leakage, thus contributing to tissue damage and inflammation (Biering et al., 2022; Luo et al., 2024). Recently, viral toxins have been suggested as therapeutic targets against emerging and re-emerging viral diseases (Biering et al., 2021; Coelho et al., 2021; Modhiran et al., 2021; Sousa et al., 2022).

Here, we tested 18 CDs with variable cavity sizes and amounts of sulfation (directly linked to the sugar ring or via a linker group) for their effect on inhibiting DENV NS1- and SARS-CoV-2 Spike-induced endothelial hyperpermeability, as well as DENV and SARS-CoV-2 replication *in vitro* and DENV infection *in vivo*.

## 2. Materials and Methods

### 2.1. CDs

A panel of 18 randomly substituted CD analogs was evaluated, including derivatives of α-cyclodextrin (ACD), β-cyclodextrin (BCD), and γ-cyclodextrin (GCD), which are composed of six, seven, and eight glucose units, respectively (Table 1). These analogs featured diverse chemical modifications, such as 2-hydroxypropyl, sulfate, and sulfobutylether substitutions. The tested compounds varied in cavity size and in the degree and type of sulfation, either directly linked to the glucose ring or introduced via a linker group. All compounds were provided by Cyclolab, Hungary.

**Table 1.** Cyclodextrin compounds and their safe concentration thresholds for anti-coagulation and cytotoxicity

| Cyclodextrin Code | Name | MW(g/mol) | Safe concentration-Anti-coagulation<br>(µg/mL) | Safe concentration-Cytotoxicity (µg/mL) |
| --- | --- | --- | --- | --- |
| CD1 | SBECD | 2179.1 | 1000 | >3000 |
| CD2 | SBEGCD | 1831.08 | 1000 | >3000 |
| CD3 | 2,3-di-Me-6-sulfate-ACD | 1753.4 | 250 | >3000 |
| CD4 | 2,3-di-Me-6-sulfate-BCD | 2045.7 | 1000 | >3000 |
| CD5 | 2,3-di-Me-6-sulfate-GCD | 2337.9 | 100 | >3000 |
| CD6 | 2,3-Ac-6-sulfo-BCD | 2437.8 | 100 | >3000 |
| CD7 | 2,3-di-OH-6-sulfo-BCD | 1849.3 | 1000 | >3000 |
| CD8 | Dexolve-10 | 2780.28 | 250 | >3000 |
| CD9 | RAMEB | 1291.8 | 1000 | >3000 |
| CD10 | HPGCD | 1576.1 | 1000 | >3000 |
| CD11 | HPBCD | 1403.4 | 1000 | >3000 |
| CD12 | Szulfopropil-ACD | 1261.1 | 1000 | >3000 |
| CD13 | Szulfopropil-BCD | 1495.4 | 1000 | >3000 |
| CD14 | Szulfopropil-GCD | 1585.4 | 1000 | >3000 |
| CD15 | 2,3-di-6-sulfo-GCD | 2113.7 | 25 | >3000 |
| CD16 | ACD-sulfate | 1902.1 | 5 | >3000 |
| CD17 | BCD-sulfate | 2360.2 | 5 | >3000 |
| CD18 | GCD-sulfate | 2318.2 | 5 | >3000 |

### 2.2. Cell lines and viruses

Human pulmonary microvascular endothelial cells (HPMEC, clone ST1-6R) were kindly provided by J.C. Kirkpatrick from Johannes Gutenberg University, Germany. These cells were cultured in Endothelial Cell Basal Medium-2 (Lonza #CC-3162), supplemented with the EGM-2 Endothelial SingleQuots Kit, following the manufacturer’s instructions. Vero cells (ATCC CCL-81), derived from African green monkey kidney epithelial cells, were maintained in Minimum Eagle Medium (MEM; Gibco) supplemented with 10% fetal bovine serum (FBS, Gibco) and 1% penicillin-streptomycin (Gibco). Both cell types were incubated at 37°C in a 5% CO_2_ atmosphere. Antiviral assays were conducted using the reference DENV strain DENV-2/S16803. Calu-3 human lung epithelial cells were obtained from the UC Berkeley Cell Culture Facility and maintained in Dulbecco Minimum Essential Medium (DMEM; Gibco) supplemented with 10% FBS and 1% penicillin-streptomycin.

### 2.3. Cell viability assay

Cytotoxicity was evaluated using the MTS (3-(4,5-dimethylthiazol-2-yl)-5-(3-carboxymethoxyphenyl)-2-(4-sulfophenyl)-2H-tetrazolium) cellular viability assay (Abcam). Briefly, HPMECs (6 × 10^4^ cells/well) were seeded in 96-well plates in 200 μL of culture medium and incubated for 72 hours. Vero cells (3 × 10^4^ cells/well) were seeded similarly and incubated for 24 hours. After the incubation period, the growth medium was removed, and the cell monolayers were treated with 2-fold dilutions of each compound (CD) ranging from 94 to 3000 µg/mL, followed by a 24-hour incubation. Cells cultured in medium only were used as the 100% viability control. Following treatment, the medium was discarded, and 200 μL of fresh medium was added to each well. Then, 200 μg/100 μL of MTS solution was added to each well, and the plates were incubated for an additional 4 hours. The optical density was measured at 490 nm. Untreated cells with medium only served as the 100% viability control. Each assay was performed in triplicate (n=3). Non-toxic concentrations were selected for use in subsequent assays.

### 2.4. Anticoagulant activity

Anticoagulant activity was assayed using the Activated Partial Thromboplastin Time (APTT XL, Pacific Homeostasis) kit, as previously described (Sousa et al., 2022). Briefly, 50 μL of each compound at 50 μg/mL was mixed with 100 μL of human plasma, then 100 μL of APTT reagent was added. The mixtures were incubated for 3 minutes at 37°C, then pre-heated calcium chloride (100 μL, 0.025 M) was added, and clotting time was recorded. PBS was used as a negative control. Heparin sodium salt (Sigma H3393–50KU), a mixture of polyanion chains with MW 6–30 kDa, was used at 0.01 ng/mL as a positive control.

### 2.5. Evaluation of CD inhibition of viral protein-induced endothelial hyperpermeability

HPMECs were cultured at 80%–90% confluency in vented 75-cm^²^ flasks (Corning^®^), detached, and resuspended in fresh culture medium. The cells were seeded onto the apical side (top chamber) of Transwell inserts at a density of 6 × 10^⁴^ cells/well, with a final volume of 300 μL per well. The inserts were then placed in a 24-well plate containing 1.5 mL of endothelial cell culture medium to represent the basolateral chamber. The Transwell plate was incubated at 37°C with 5% CO_2_ for 3 days, during which 50% of the culture medium in the apical (top) chamber was replaced daily, and the basolateral medium was replaced every 48 hours. Cells were cultured until their trans-endothelial electrical resistance (TEER) reached between 150 and 180 Ω, indicating full cell confluency. Once confluency was achieved, recombinant DENV NS1 protein (10 μg/mL), SARS-CoV-2 Spike protein (10 μg/mL), or a mixture of the proteins with the CDs were added to the apical side of the Transwell inserts (top chamber, 300 μL). TEER measurements were taken every 2 hours using an Epithelial Volt-Ohm Meter (EVOM^2^) with chopstick electrodes (World Precision Instruments). Endothelial permeability was expressed as relative TEER, calculated as the ratio of resistance values (Ω) according to the formula:

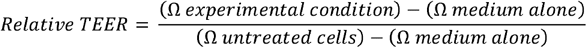

Maximum hyperpermeability values were recorded at 6- and 24-hours post-treatment for DENV NS1 and SARS-CoV-2 Spike proteins, respectively, and represented as the area under the curve.

### 2.6. Evaluation of antiviral activity of cyclodextrins

To evaluate the antiviral activity of CDs against DENV2, Vero cells (3 × 10^⁴^ cells/well) were plated and infected with an MOI of 1 and 0.1. The cells were treated with CDs simultaneously with the infection and incubated for 2 hours. After incubation, the virus was removed, and fresh medium with a new treatment of CDs was added. The cells were then incubated for an additional 24-48 hours. After this period, supernatants were collected to assess the antiviral effect. Viral titers in the supernatants were determined using a focus-forming assay. In parallel, the virucidal effect of the compounds was tested by pre-incubating the virus with the CDs before infecting the monolayers. Viral titers were again determined using a focus-forming assay. Celgosivir (20 µg/mL) was used as a positive control (Rathore et al., 2011; Watanabe et al., 2012).

To assess the antiviral effect of CDs against SARS-CoV-2, Calu-3 cells (1 × 10^⁵^ cells/well) were seeded in 24-well plates and incubated for 48 hours. The plates were then transferred into a biosafety level 3 (BSL-3) facility and infected with SARS-CoV-2 at MOI of 0.05 in a total volume of 300 µL. The indicated concentrations of CDs or reference compounds were added directly to the inoculum during infection. After a 30-minute incubation, the inoculum was removed, the cells were washed with 1X PBS, and 1 mL of fresh D10 medium containing the respective treatments was added. At 24 hours post-infection, culture supernatants were harvested and stored at –80°C. Viral titers in supernatants were determined using the tissue culture infectious dose 50% (TCID_50_) assay on Vero cells and expressed as infectious units per milliliter as previously described (Biering et al., 2022; Biering et al., 2021).

### 2.7. Evaluation of the therapeutic potential of CD1 in a DENV mouse model

Six-to eight-week-old female C57BL/6 interferon (IFN) α/β receptor knockout (*Ifnar*^-/-^) mice bred in the UC Berkeley Animal Facility were infected intravenously with 3.6 × 10^6^ PFU/mL of DENV2 D220 (Orozco et al., 2012). Starting on the day of infection (day 0) through day 5, each mouse was administered 10 mg/kg of CD1 diluted in PBS or PBS alone as a negative control twice per day via intraperitoneal inoculation. Mice were observed daily for morbidity and mortality over the next 14 days. Morbidity of mice was assessed utilizing a standardized 1-5 scoring system as follows: 1 = no signs of lethargy, mice are considered healthy; 2 = mild signs of lethargy and fur ruffling; 3 = hunched posture, further fur ruffling, failure to groom, and intermediate level of lethargy; 4 = hunched posture with severe lethargy and limited mobility, while still being able to cross the cage upon stimulation; and 5 = moribund with limited to no mobility and inability to reach food or water (euthanized). Mice were also weighed to measure weight loss over days post-infection. All experimental procedures involving animals were approved by the Animal Care and Use Committee (ACUC) of the University of California, Berkeley.

### 2.8. Statistical analysis

Statistical comparisons of test groups for *in vitro* and *in vivo* assays were analyzed by Students t-test, Wilcoxon Mann-Whitney test, or ANOVA followed by multiple comparison tests, as appropriate. For *in vitro* assays, compounds were tested in 3 separate experiments, with each concentration run in duplicate, and the results were averaged to yield mean and standard deviation. Significant differences were considered p<0.05. Kaplan-Meier survival curves were compared using a non-parametric log-rank (Mantel-Cox) test. All analyses were performed in GraphPad Prism.

## 3. Results

### 3.1. CDs displayed no significant cytotoxicity or anticoagulant activity at the doses tested

First, we assessed the cytotoxicity of serial dilutions of each CD derivative (Figure 1) and compound using the MTS assay. This initial step was particularly important given that sulfated saccharides, due to their polyanionic nature, can influence both cell viability and nonspecific interactions with cellular membranes and plasma proteins. To evaluate cytotoxicity, Calu-3 and HPMECs cells were exposed to 2-fold serial dilutions of each CD, ranging from 93.4 to 3000⍰µg/mL, and viability was measured after 24 hours. The highest concentration that did not significantly reduce viability compared to untreated controls was defined as the safe concentration (Table 1, Supplementary Figure 1). Moreover, the potential anticoagulant activity of CDs must be considered, as such effects could exacerbate vascular leak, especially in viral infections where bleeding is a common and clinically relevant complication. To evaluate this, *in vitro* anticoagulant activity was assessed using human plasma in the activated partial thromboplastin time (APTT) assay. Coagulation times were compared to plasma treated with PBS alone. Several CDs exhibited significant anticoagulant activity and were further diluted until no significant prolongation of clotting time was observed (p > 0.05; Supplementary Figure 2). Thus, based on the cytotoxicity and coagulation findings, we selected concentrations for each CD that exhibited no direct harmful effects on HPMECs for use in subsequent antiviral assays (Table 1).

**Figure 1.**
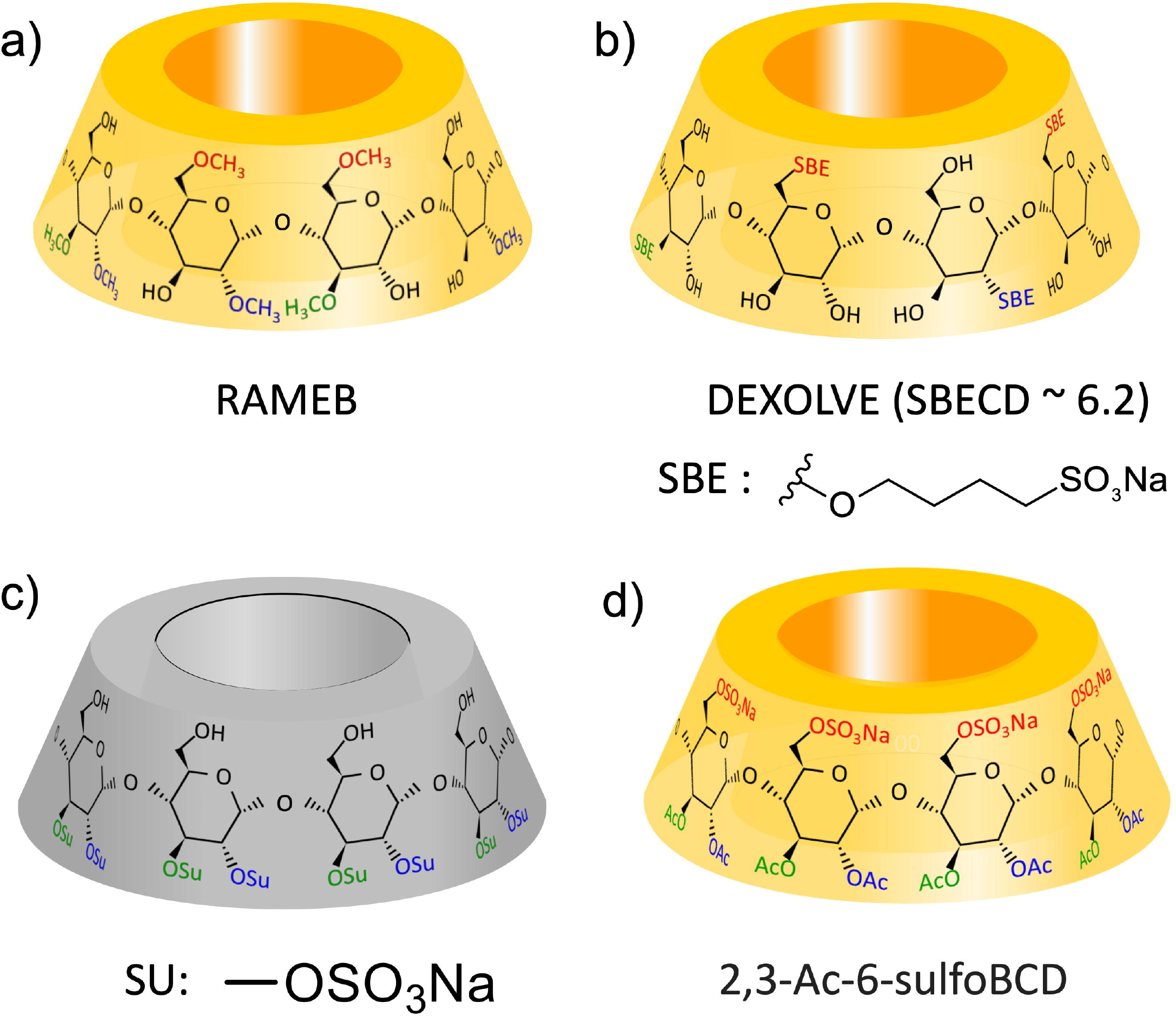
Representative chemical structures of selected cyclodextrins (CDs) used in this study. Structures of four representative CD derivatives evaluated for antiviral activity. **(a)** RAMEB (CD-X) is a randomly methylated β-cyclodextrin. **(b)** Dexolve (SBECD [CD-Y], degree of substitution ∼6.2) is a sulfobutylether-β-cyclodextrin widely used in pharmaceutical formulations. **(c)** The sulfonate substituent (SU: -OSO_₃_Na) represents direct sulfation of hydroxyl groups (CD-Z). **(d)** 2,3-Ac-6-sulfo-β-cyclodextrin (2,3-Ac-6-sulfoBCD; CD-A) is a trisubstituted CD bearing acetyl groups at positions 2 and 3 and a sulfonate group at position 6 of each glucose unit. These structural variations modulate cavity polarity, solubility, and host-target interactions relevant to biological activity.

### 3.2. CDs inhibited flavivirus NS1-triggered endothelial hyperpermeability

Soluble NS1 is a complex virulence factor present in the blood of patients infected with flaviviruses. A key mechanism by which NS1 causes barrier dysfunction is through its direct interaction with endothelial cells. This interaction triggers signaling pathways that lead to the disruption of endothelial barriers, causing increased permeability in endothelial cell monolayers cultured in Transwells (Beatty et al., 2015; Puerta-Guardo et al., 2016). We hypothesized that CDs could inhibit the capacity of DENV NS1 to trigger endothelial hyperpermeability. To evaluate this, we utilized a TEER assay to monitor barrier resistance across a monolayer of HPMECs seeded in the apical chamber of a Transwell. As previously observed, we found that DENV NS1 triggers endothelial hyperpermeability of HPMEC (Fig. 2a). In contrast, treating cells with NS1 in the presence of 13 of the 18 CDs tested reduced the capacity of NS1 to induce endothelial hyperpermeability (Fig. 2a). These data demonstrate the potential of CDs to prevent hyperpermeability induced by DENV NS1.

**Figure 2.**
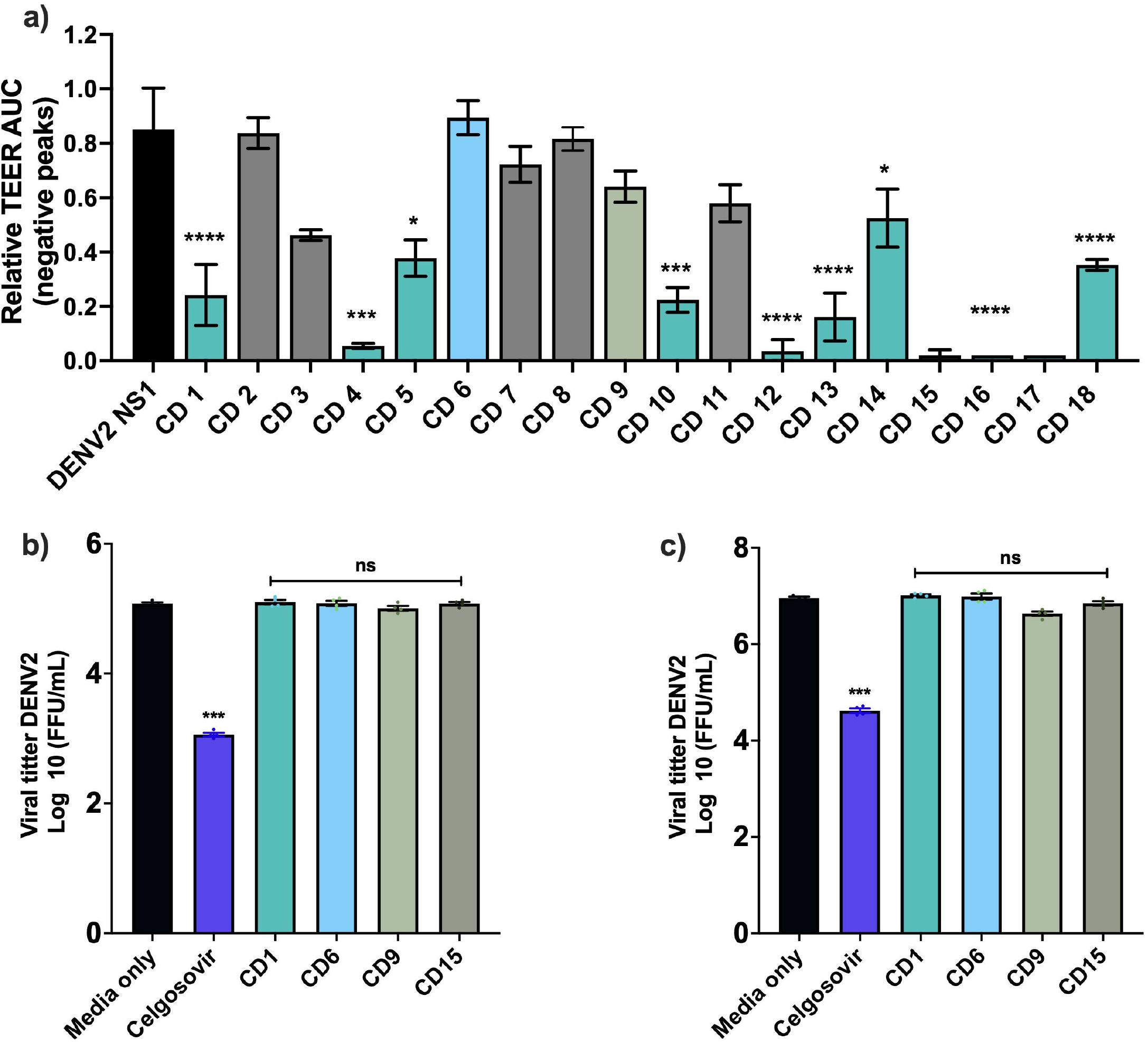
Inhibition of DENV2 NS1-induced endothelial hyperpermeability and antiviral activity against DENV2 of modified CDs. **(a)** Select CDs were evaluated for their ability to prevent NS1-mediated hyperpermeability. Trans-endothelial electrical resistance (TEER) was monitored between 2- and 10-hours post-treatment. The area under the curve (AUC) of TEER measurements was calculated; higher AUC values indicate greater permeability, while lower values reflect a protective effect, maintaining barrier integrity. Data are presented as mean ± standard error of the mean (SEM, n=4). Statistical significance was determined by one-way ANOVA followed by Dunnett’s post-hoc test, comparing each treatment to the DENV2 NS1-only condition at 10⍰μg/mL. **(b-c)** Evaluation of the antiviral activity of selected cyclodextrins (CD1, CD6, CD9, CD15) against DENV2 (strain 16803) in Vero cells. Cells (3 × 10^⁴^/well) were infected at a multiplicity of infection (MOI) of 0.1 and treated with the indicated compounds or the α-glucosidase inhibitor Celgosivir 20ug/mL (positive control). Treatments were administered both during and after infection. Viral titers in the supernatants were quantified by focus-forming assay on Vero cells at 48 hours (c) post-infection. Data are expressed as log_10_ FFU/mL ± SEM (n = 4). Statistical analysis was performed using one-way ANOVA followed by Dunnett’s post-hoc test vs. NS1-treated group. ^*^p⍰<⍰0.05, ^***^p⍰<⍰0.001, ^****^p⍰<⍰0.0001; gray bars: ns (not significant).

### 3.3. Select CDs did not inhibit DENV infection in cell culture

We then evaluated the antiviral efficacy of select CDs *in vitro* using a focus-forming assay to assess viral titers from supernatants of DENV2-infected Vero cells. Of the 18 CD derivatives screened in the TEER assay, we selected 4 compounds for follow-up infection studies based on their differential activity profiles. CD1 was included primarily due to prior reports in the literature supporting its ability to modulate host membrane composition and its known safety profile (Pardeshi et al., 2023). CD15 was selected for its activity at lower concentrations compared to other active compounds identified in this screen and its structural similarity to other sulfated CDs tested in the same panel. In contrast, CD5 and CD11 did not significantly inhibit NS1-induced barrier disruption in TEER assays and were used as negative controls to confirm the specificity of the protective effects observed with the active compounds. Cells were treated with various concentrations of the compounds simultaneously with viral infection. No inhibitory effect was observed for any of the CDs tested at the concentrations used (Fig. 2b and 2c). As a positive control, Celgosivir, an α-glucosidase I inhibitor, demonstrated antiviral activity under both co-treatment and post-infection conditions (Fig. 2b, 2c) (Watanabe et al., 2012). These results indicate that the tested CDs did not impair DENV replication under the conditions evaluated.

### 3.4. CDs reduced SARS-CoV-2 infection in vitro and Spike-induced endothelial hyperpermeability

SARS-CoV-2 infection is strongly associated with endothelial dysfunction, a process exacerbated by the viral Spike protein (Biering et al., 2022). To evaluate the potential of CDs as protective agents against endothelial dysfunction associated with COVID-19, we first assessed their ability to preserve endothelial barrier integrity in the presence of recombinant Spike protein. The 4 compounds (CD1, CD15, CD5, and CD11) were the same subset selected for next assays based on their differential activity profiles observed in the NS1-induced permeability screen, allowing us to directly compare their performance across distinct viral triggers of endothelial disruption. Using TEER measurements in HPMEC monolayers, we found that 2 out of 4 CDs tested (CD1, CD15) significantly inhibited Spike-induced endothelial hyperpermeability (Fig. 3a-b). We then investigated whether the CDs also affected SARS-CoV-2 replication. In Calu-3 epithelial cells, 13 out of the 18 CDs tested (including CD1 and CD15) significantly reduced viral titers, indicating antiviral activity (Fig. 3c). Together, these findings suggest that certain CDs offer dual functionality by protecting the endothelial barrier and reducing viral replication, supporting their potential as therapeutic agents against SARS-CoV-2 infection and its associated vascular complications.

**Figure 3.**
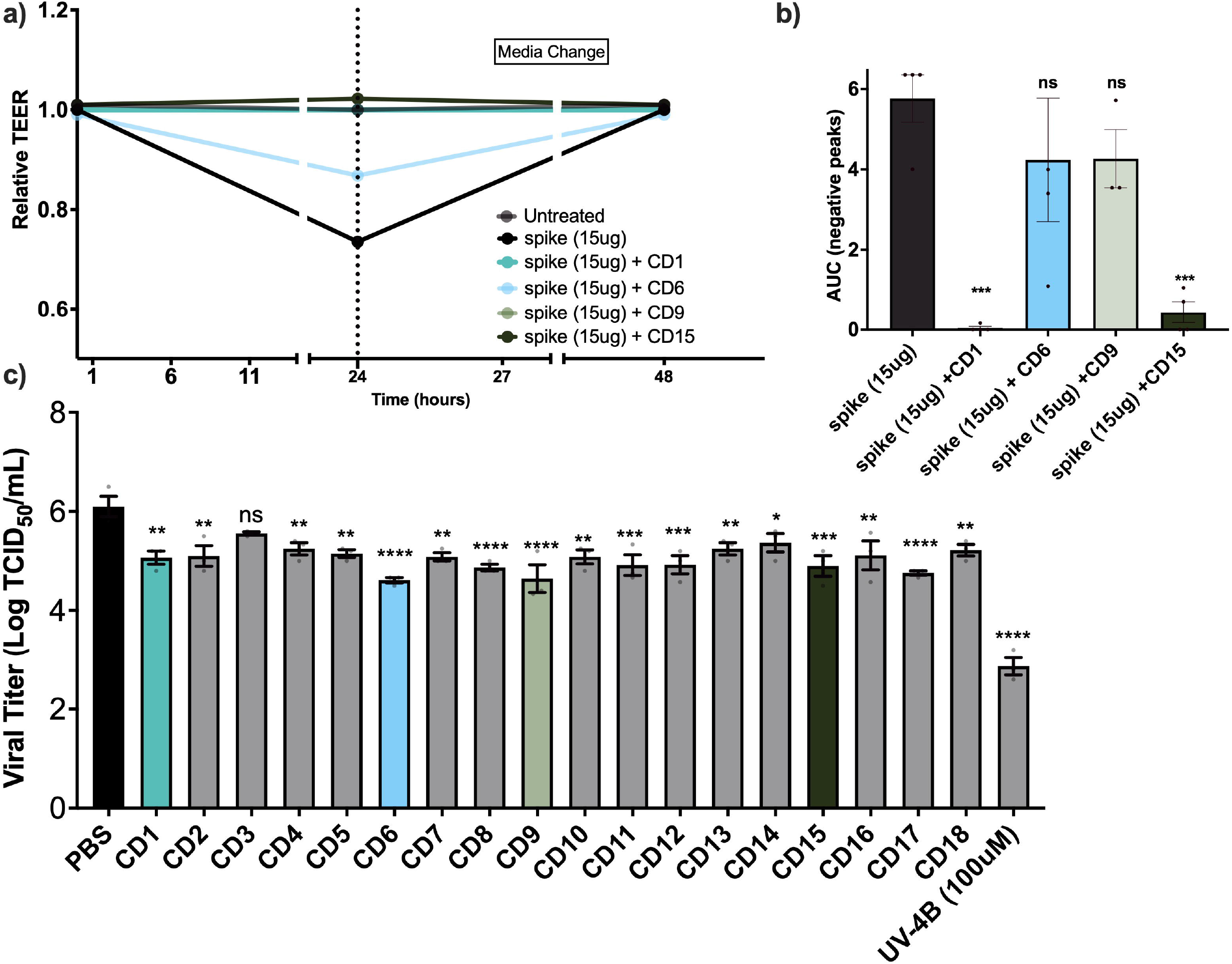
CDs protect against SARS-CoV-2 Spike-induced endothelial dysfunction and reduce viral infection. **(a)** TEER was measured in HPMEC monolayers treated with 15illµg/mL recombinant SARS-CoV-2 Spike protein, alone or in combination with selected cyclodextrins (CD1, CD6, CD9, or CD15). TEER values were recorded at 0, 24, and 48 hours. Medium was changed at 24 hours (dotted line). CD1 and CD15 significantly reversed Spike-induced endothelial hyperpermeability, as reflected by higher TEER values compared to Spike-only treatment. **(b)** Quantification of endothelial barrier disruption by calculating the area under the curve (AUC) of negative TEER peaks. Treatment with CD1 (^***^p⍰<⍰0.001) or CD15 (^**^p⍰<⍰0.01) significantly reduced AUC compared to Spike-only, while CD6 and CD9 showed no significant effect (ns = not significant). Data represent mean ± SEM from 3 independent experiments. Statistical analysis was performed using one-way ANOVA with Dunnett’s multiple comparisons test. **(c)** Antiviral activity of CDs against SARS-CoV-2. Calu-3 cells were infected at an MOIillof 0.05 in the presence of CDs or control compounds. Treatments were applied during inoculation and maintained post-infection. Supernatants were collected at 24 hours, and viral titers quantified by TCID_50_ on Vero-CCL81 cells. Treatment with 13 CDs significantly reduced viral replication, indicating antiviral potential. Data are shown as mean ± SEM (n⍰=⍰3–4). Statistical analysis was performed using one-way ANOVA followed by Dunnett’s post-hoc test. ^*^p⍰<⍰0.05; ^**^p⍰<⍰0.01; ^***^p⍰<⍰0.001; ^****^p⍰<⍰0.0001; ns = not significant.

### 3.5. CD1 protected mice from lethal DENV challenge

Among the panel of CDs evaluated, CD1 emerged as the most promising candidate for *in vivo* testing based on its broad-spectrum protective effects against viral toxin-induced endothelial dysfunction and its status as an FDA-approved compound. As shown in the Venn diagram (Fig. 4), CD1 was one of only two compounds that significantly preserved endothelial barrier integrity inhibiting both DENV2 NS1- and SARS-CoV-2 Spike-induced hyperpermeability, as measured by TEER assays. This dual specificity highlights its potential as a therapeutic capable of mitigating vascular leak, a hallmark of severe dengue and COVID-19 disease. CD1 also showed antiviral activity against SARS-CoV-2 infection.

**Figure 4.**
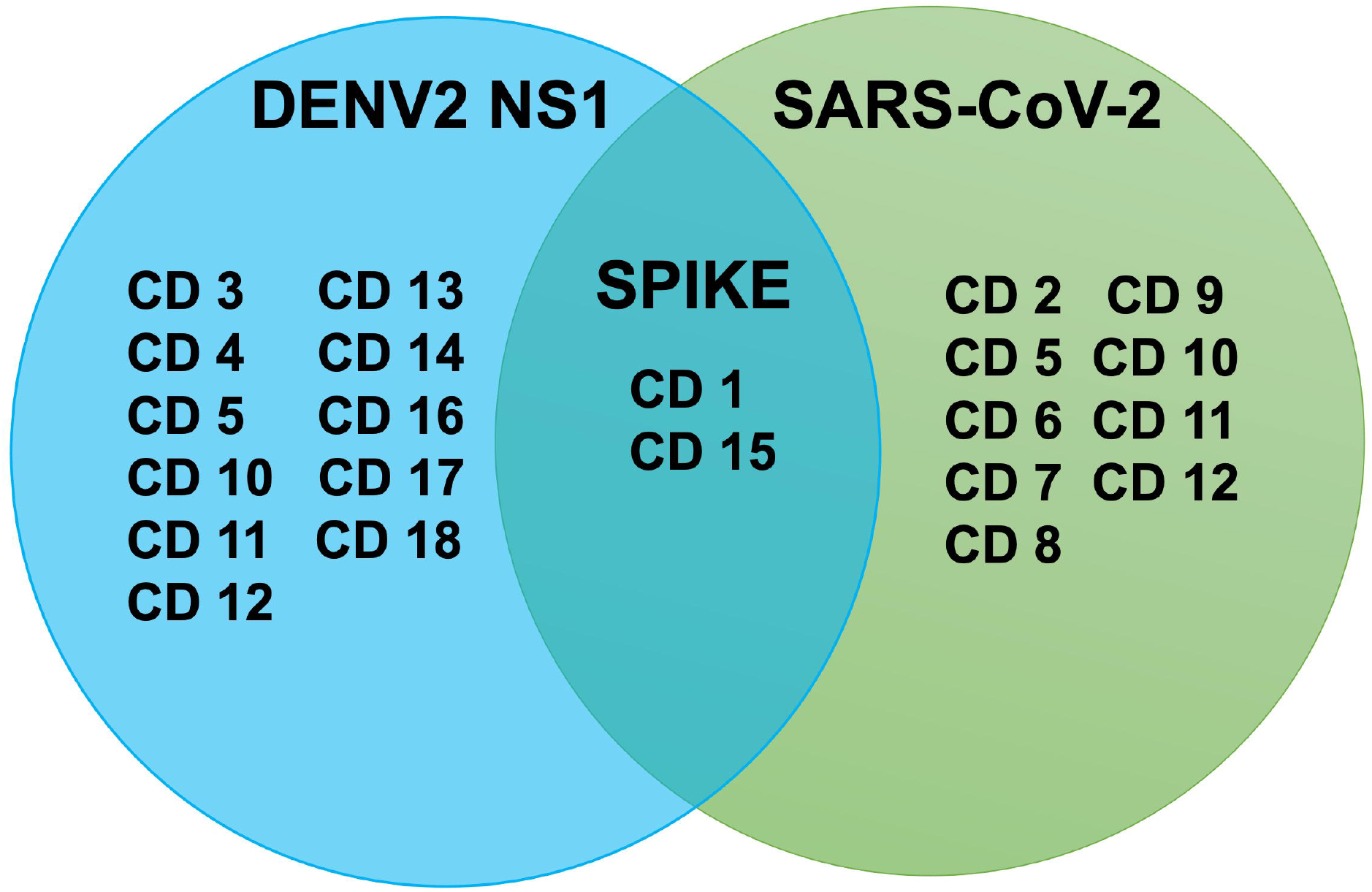
Overlap of CDs that protect against SARS-CoV-2 infection or endothelial dysfunction induced by DENV2 NS1 and SARS-CoV-2 Spike proteins. The Venn diagram illustrates the modified CDs that significantly reduced SARS-CoV-2 infection (right circle) or reversed disruption of endothelial barrier integrity in response to DENV2 NS1 (left circle) or SARS-CoV-2 Spike (overlap; CD1, CD6, CD9, CD15 assessed by TEER assay). CDs located in the overlap zone (CD1 and CD15) exhibited protective effects in all 3 models, indicating their broad-spectrum potential to mitigate virus infection and endothelial hyperpermeability.

To further evaluate the protective efficacy of CD1 in vivo, we used a DENV2 mouse-adapted virus (D220) in an interferon *α*/β receptor-deficient (*Ifnar*^-/-^) mouse model, as detailed previously (Biering et al., 2021; Orozco et al., 2012). Mice received a lethal challenge of DENV2 D220 intravenously (IV) on day 0 and daily intraperitoneal (IP) injections of the indicated dose of CD1 from days 0 to 4 post-infection (5 IP infections in total) (Fig. 5). We observed protection from virus-induced morbidity and mortality over 14 days, where 33% of mice receiving 5 doses of 10 mg/kg were protected from mortality in comparison to the vehicle control (Fig. 3b). In addition, mice began recovering weight and displayed improved morbidity scores ∼8 days post-infection compared to controls (Fig. 3c-d). These results demonstrate the potential of CD1 treatment to protect in a mouse model of lethal DENV vascular leak syndrome.

**Figure 5.**
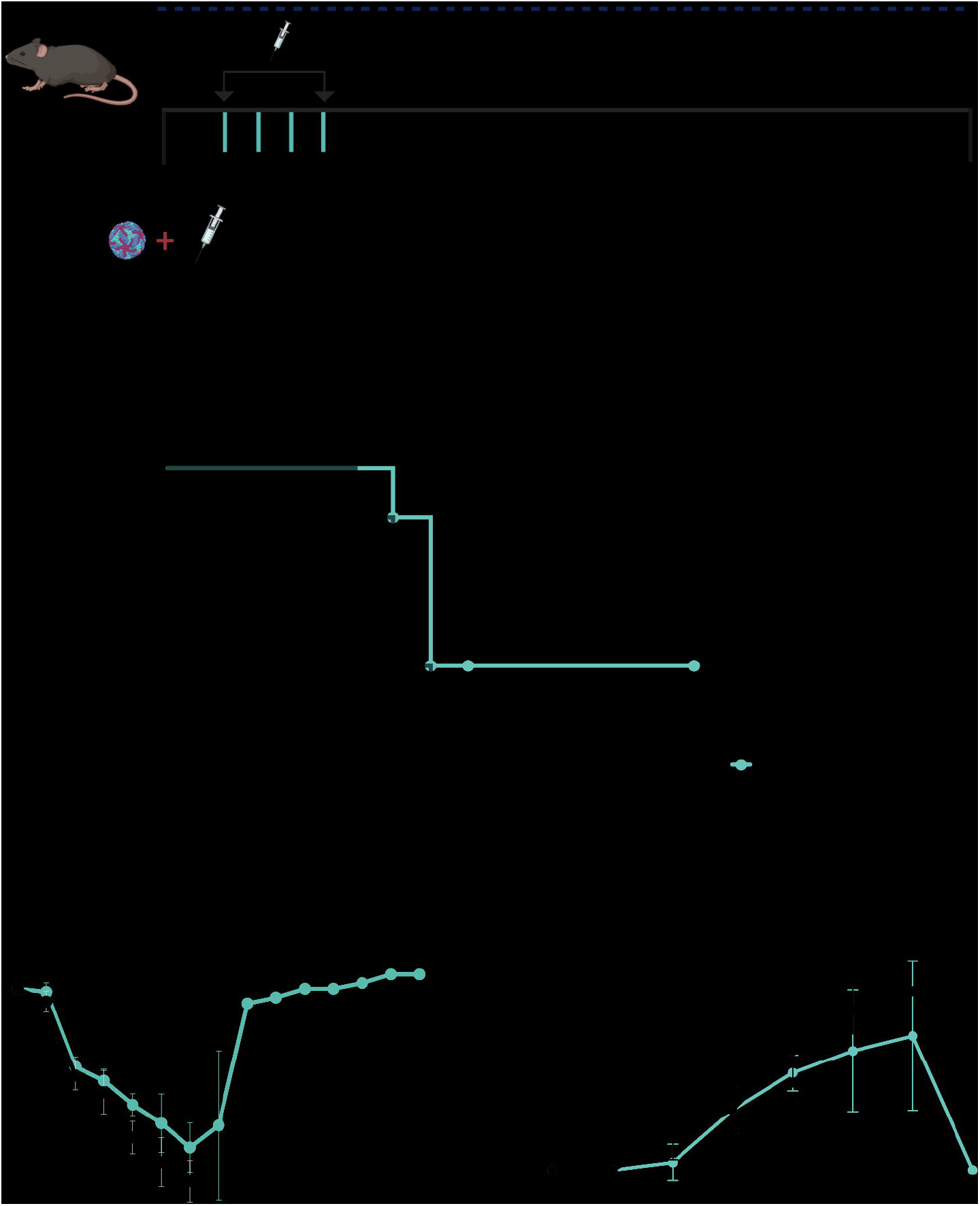
*In vivo* protection against lethal DENV infection by CD1 treatment. **(a)** To evaluate the protective efficacy of CD1 in vivo, C57BL/6 Ifnar^−/−^ mice (n=6 per group) were infected intravenously (IV) on day 0 with 3.6 × 10^⁶^ FFU/mL of DENV2 (strain D220) in a volume of 100 μL. Mice received daily intraperitoneal (IP) injections of CD1 (10 mg/kg) or PBS vehicle from day 0 through day 4 post-infection (5 total doses). Mice were monitored daily for signs of disease through day 14. **(b)** Kaplan–Meier survival curves comparing CD1-treated and PBS-treated groups. Statistical significance was determined by Mantel–Cox log-rank test; CD1 treatment significantly improved survival (p = 0.0468). **(c)** Weight and **(d)** morbidity scores were recorded daily based on a clinical scoring scale from 1 (healthy) to 5 (moribund or dead). CD1-treated mice showed a trend toward reduced morbidity compared to controls. Data represent mean ± SEM (n = 6 per group). This figure was partially created in biorender.

## 4. Discussion

CDs have emerged as promising antiviral agents with dual functionality: direct antiviral activity and protection of host barrier integrity (Braga et al., 2021; Jones et al., 2020). In this study, we demonstrate that specific CD analogues effectively inhibit endothelial hyperpermeability induced by viral proteins from DENV (NS1) and SARS-CoV-2 (Spike), without compromising host cell viability. Selected CDs also reduced SARS-CoV-2 replication *in vitro*, and CD1 conferred partial protection against DENV-induced morbidity and mortality *in vivo*. CDs have a low toxicity in humans and are currently used as excipients (Braga et al., 2021; Hastings et al., 2022). These findings highlight the therapeutic potential of CDs in mitigating both viral replication and virus-driven vascular pathology, offering a novel, virus- and host-directed approach for mitigating severe complications associated with dengue and COVID-19.

During viral infection, barrier disruption is a critical pathological event that can facilitate viral dissemination (Puerta-Guardo et al., 2024, Pahmeier et al., 2025). Endothelial and epithelial barriers, including the glycocalyx, play critical roles in maintaining vascular integrity. Specific viral proteins have been shown to target these barriers directly, impairing their structural and functional integrity. Dengue NS1 protein and the SARS-CoV-2 Spike protein are recognized not only for their roles in viral replication (Glasner et al., 2018) and host entry (Yu et al., 2023), respectively, but also as major contributors to virus-induced barrier dysfunction.

DENV NS1 has been extensively characterized as a key mediator of endothelial hyperpermeability during dengue pathogenesis, disrupting the endothelial glycocalyx and intercellular junction complex integrity (Biering et al., 2021; Puerta-Guardo et al., 2019, 2016). DENV NS1 exerts diverse pathogenic effects on endothelial cells, including the promotion of endothelial dysfunction, induction of vascular leak, and activation of inflammatory responses in immune cells (Modhiran et al., 2017, 2015, Beatty et al., 2015), believed to contribute to the hallmark manifestations of severe dengue, such as plasma leakage syndrome. Similarly, the SARS-CoV-2 Spike protein, while primarily facilitating viral entry into epithelial cells via the receptor ACE2, has also been shown to directly impair epithelial and endothelial function independently of ACE2 (Biering et al., 2022, Panigrahi et al., 2021). In epithelial cells, exposure to Spike protein leads to downregulation of ACE2 expression, activation of apoptotic pathways, and disruption of vascular integrity, contributing to inflammation and coagulopathies and exacerbating vascular leakage and tissue damage (Biering et al., 2022; Hussein et al., 2024; Panigrahi et al., 2021).

Here, we demonstrated that multiple CDs significantly inhibited endothelial hyperpermeability induced by DENV NS1 *in vitro* without impacting DENV replication in cells. *In vivo*, CD1 provided partial protection against virus-induced mortality and improved morbidity scores in a murine model of DENV vascular leak syndrome. These findings suggest that the protective effects of CD1 may be mediated through interference with NS1-driven endothelial dysfunction and disease severity rather than direct antiviral activity. This distinction underscores the potential utility of CD-based interventions focused on mitigating virus-induced vascular pathology, an approach that could complement traditional strategies targeting viral replication. Furthermore, CD1 offers regulatory and practical advantages, as it is an FDA-approved compound commercially available in the United States, and it is recognized for its high purity standards -- factors that make it well-suited for translational research and preclinical and clinical evaluation.

Regarding SARS-CoV-2, our results are in line with current knowledge about the association between the Spike protein and endothelial dysfunction (Biering et al., 2022; Luo et al., 2024; Panigrahi et al., 2021). Treatment with selected CDs not only preserved endothelial barrier integrity against Spike-induced hyperpermeability but also significantly reduced viral replication *in vitro*. These dual protective effects reinforce the concept that CDs can act at multiple stages of the viral pathogenic process.

Beyond their well-known role in enhancing drug solubility and stability (Kovacs et al., 2022), CDs have emerged as promising antiviral agents. Recent studies have demonstrated that native and chemically modified CDs, including hydroxypropyl-β-cyclodextrin (HPβCD) and sulfobutylether-β-cyclodextrin (SBEβCD), can interfere with viral infection by modulating host cell membrane composition and depleting cholesterol, which is critical for viral entry and fusion (Jones et al., 2020; Senti et al., 2013). Notably, HPβCD has been shown to inhibit SARS-CoV-2 replication in Calu-3 pulmonary cells, impair viral fusion via cholesterol depletion, and exert prophylactic effects in nasal epithelium models in vivo (Almeida et al., 2023; Jones et al., 2020; Raïch-Regué et al., 2023). These findings are consistent with our observations and suggest that the antiviral activity of CDs may stem from both direct inhibition of viral replication and modulation of viral protein-host membrane interactions.

Of particular relevance, SBEβCD—the active compound in CD1—is a chemically modified β-cyclodextrin with a well-characterized safety profile. It is already approved for intravenous administration in several pharmaceutical formulations, including that of the antiviral agent against SARS-CoV-2, remdesivir (Veklury^®^). Structural modifications in SBEβCD enhance its aqueous solubility and substantially reduce nephrotoxicity and hemolytic activity compared to native β-cyclodextrins, making it highly suitable for systemic therapeutic applications (Pardeshi et al., 2023).

The findings that CDs are effective against both the SARS-CoV-2 Spike protein and SARS-CoV-2 infection are consistent with the biological properties of Spike. As a structural component of the virion, Spike is critical for host cell entry via ACE2 and can also contribute to epithelial barrier disruption during infection (Li et al., 2023). Additionally, Spike can be shed from infected cells into circulation as a soluble protein, independently inducing endothelial dysfunction and inflammatory responses and contributing to vascular pathology even in the absence of infectious virus (Biering et al., 2022; Luo et al., 2024). In contrast, DENV NS1 is a non-structural protein that is not integral to the DENV virion. NS1 contributes to dengue pathogenesis by enhancing viral replication intracellularly and, once secreted, promoting endothelial dysfunction and modulating immune responses in the bloodstream (Scaturro et al., 2015, Beatty et al., 2015; N. Modhiran et al., 2015; Puerta-Guardo et al., 2022). Thus, it is plausible that CDs target the endothelial dysfunction triggered by NS1 without necessarily having an antiviral effect on DENV replication.

Given their extracellular localization, both Spike and NS1 are accessible to therapeutic agents without requiring intracellular delivery. However, the nature of their association with host or viral surfaces differs. Targeting Spike involves interactions with the virion surface or with shed Spike protein circulating in the bloodstream, whereas targeting NS1 requires engagement with its secreted or cell-associated forms, which are implicated in vascular and immune dysfunction. The therapeutic efficacy of CD1 in reducing DENV-induced morbidity and mortality in mice further underscores the critical role of secreted NS1 in disease progression. These distinctions emphasize the importance of considering the extracellular presence and functional role of viral proteins when designing antiviral strategies. Given their biocompatibility, versatility, and safety profile, CDs represent promising candidates for the development of targeted extracellular antivirals capable of mitigating virus-induced barrier dysfunction and systemic pathology across different viral infections.

Indeed, CDs have shown activity against a wide range of viruses, including enveloped and non-enveloped viruses. Their broad-spectrum antiviral properties make them attractive candidates for the development of novel antiviral therapies that can target multiple viral infections simultaneously. Our study evaluated 18 CD analogues for their potential to mitigate barrier dysfunction, a key pathophysiological feature associated with DENV and SARS-CoV-2 infections. Our findings reveal the ability of CDs to target endothelial dysfunction induced by DENV NS1 and SARS-CoV-2 spike proteins *in vitro*, underscoring their therapeutic potential in combating vascular leakage and associated complications. Additionally, selected CDs demonstrated promising antiviral activity against SARS-CoV-2 replication *in vitro*, suggesting a dual mechanism of action against viral infection and its associated endothelial dysfunction. Furthermore, our *in vivo* studies highlight the therapeutic efficacy of the most active compound (CD1) in mitigating DENV-induced morbidity and mortality, emphasizing the translational potential of CDs as novel therapeutic agents for the treatment of dengue and COVID-19, especially since certain CDs (e.g., CD1) are already FDA-approved. Overall, our findings provide valuable insights into the potential therapeutic utility of CDs in targeting viral pathogenesis and associated complications, paving the way for further exploration and development of CD-based therapies for viral infections.

## ACKNOWLEDGMENTS

This work was supported by NIAID/NIH grants R01 AI24493 (E.H.) and R01 AI168003 (E.H.), R21 AI146464 Supplement (E.H.), and a Fast Grant from Emergent Ventures (E.H.). S.B.B. was supported in part as an Open Philanthropy Awardee of the Life Sciences Research Foundation.

## Authors contributions statement

P.H.C. Conceptualization; Data curation; Formal analysis; Methodology; Investigation; Writing - original draft; Writing - review & editing

E.V.J.P Conceptualization; Data curation; Formal analysis; Methodology; Investigation; Writing - original draft; Writing - review & editing

F.T.G.S Conceptualization; Data curation; Formal analysis; Methodology; Investigation

S.B.B. Conceptualization; Data curation; Methodology; Investigation; Formal analysis; Writing - review & editing

T.S.P. Methodology; Investigation

S.B. Methodology; Investigation

J.P. Resources; Writing - review & editing

S.S. Resources

P.L. Conceptualization; Writing - review & editing

T.S. Conceptualization; Resources; Writing - review & editing

R.B Conceptualization; Supervision; Validation; Funding acquisition; Project administration; Writing - original draft; Writing - review & editing

E.H. Conceptualization; Funding acquisition; Project administration; Resources; Supervision; Writing - review & editing

## FIGURE LEGENDS

**Figure S1:**
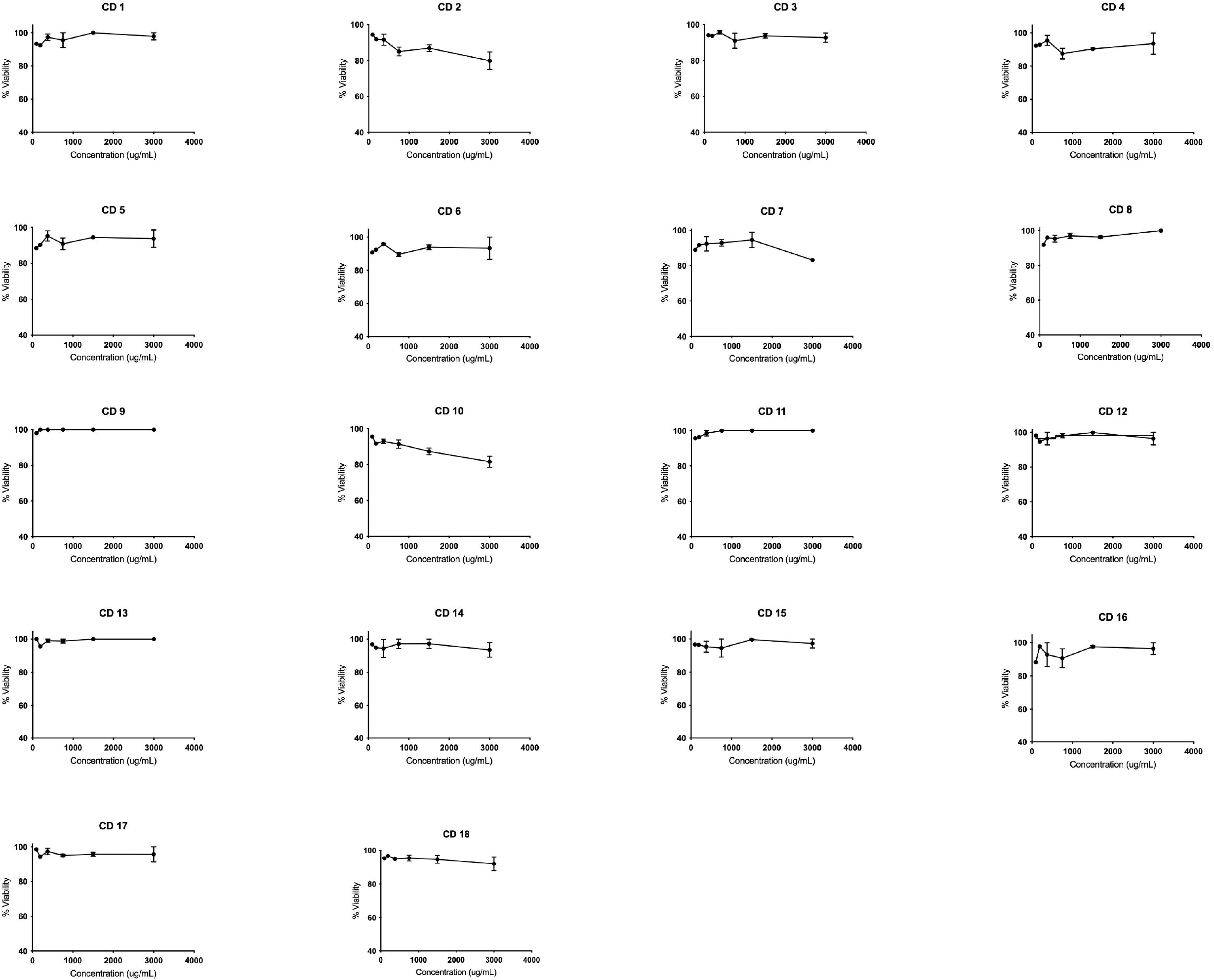
Representative cytotoxicity profiles of CDs (CD1–CD18) in HPMECs. Cell viability was assessed using the MTS assay after a 24-hour treatment with increasing concentrations of each CDs (93.75–3000 µg/mL), followed by incubation with MTS reagent. Viability was calculated relative to untreated control cells (set at 100%). Data represent the mean ± SD of three independent experiments (n = 3). No significant cytotoxicity was observed at any of the tested concentrations. Graphs shown correspond to HPMECs as a representative cell type; comparable results were obtained in other cell lines tested.

**Figure S2.**
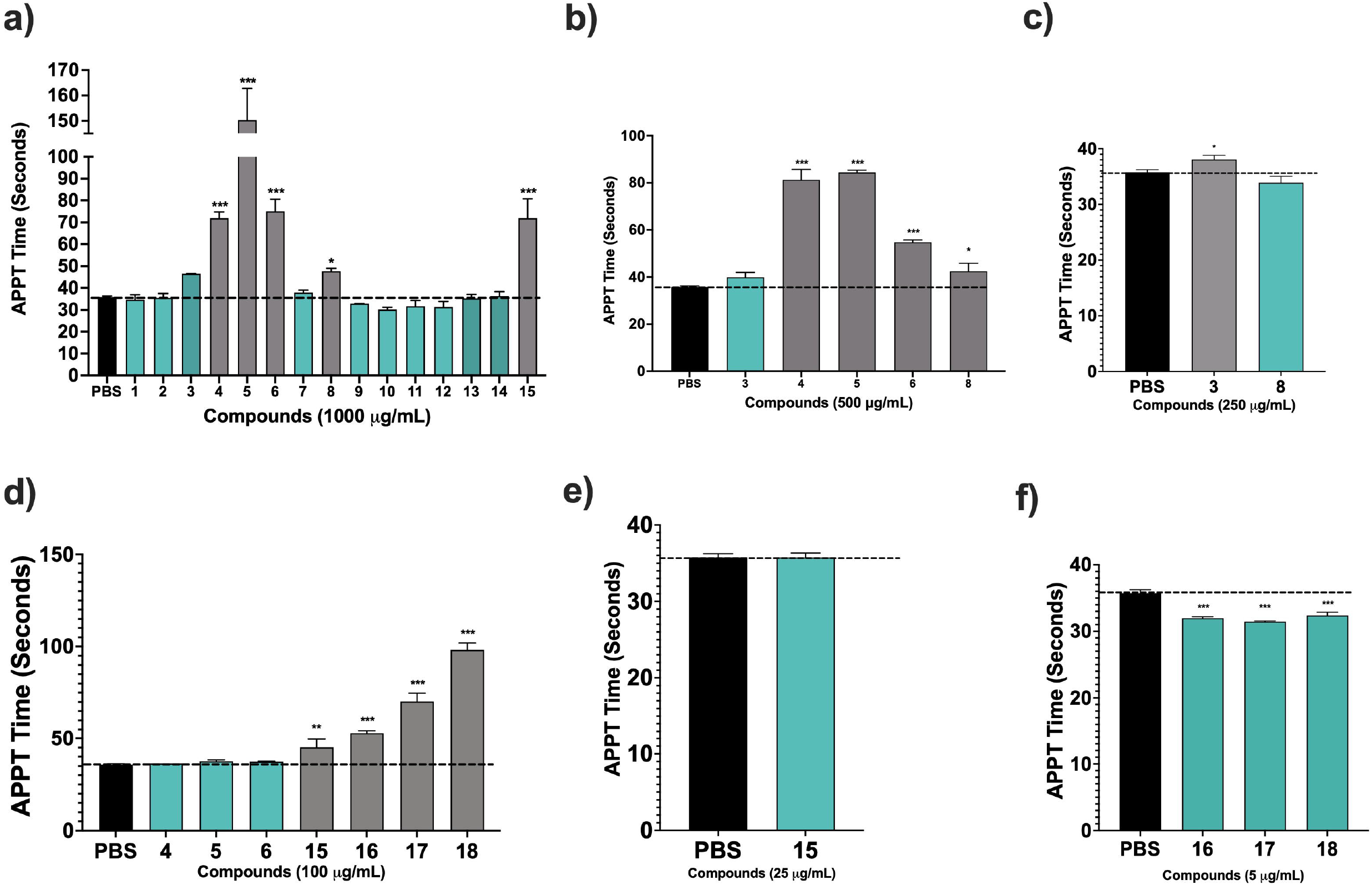
Anticoagulant activity of CDs *in vitro*. **(a-f)** The potential anticoagulant effects of CDs were evaluated using the activated partial thromboplastin time (APTT) assay in normal human plasma at concentrations of 1000⍰µg/mL, 500⍰µg/mL, 250⍰µg/mL, 100⍰µg/mL, 25⍰µg/mL, and 15⍰µg/mL, as indicated. Statistical comparisons between treatment groups and the PBS control were performed using one-way ANOVA followed by Dunnett’s multiple comparisons. Differences were considered statistically significant at p < 0.05. Green bars indicate concentrations at which no statistically significant anticoagulant activity was observed compared to PBS (ns = not significant), suggesting these doses are within a safe range.

## REFERENCES

Alboni, S., Secco, V., Papotti, B., Vilella, A., Adorni, M.P., Zimetti, F., Schaeffer, L., Tascedda, F., Zoli, M., Leblanc, P., Villa, E., 2023. Hydroxypropyl-β-Cyclodextrin Depletes Membrane Cholesterol and Inhibits SARS-CoV-2 Entry into HEK293T-ACEhi Cells. Pathogens 12(5), 647. 10.3390/pathogens12050647

Almeida, B., Domingues, C., Mascarenhas-Melo, F., Silva, I., Jarak, I., Veiga, F., Figueiras, A., 2023. The Role of Cyclodextrins in COVID-19 Therapy-A Literature Review. Int J Mol Sci 24 (3), 2974. 10.3390/ijms24032974

Beatty, P.R., Puerta-Guardo, H., Killingbeck, S.S., Glasner, D.R., Hopkins, K., Harris, E., 2015. Dengue virus NS1 triggers endothelial permeability and vascular leak that is prevented by NS1 vaccination. Sci Transl Med 7 (304), 304ra141. 10.1126/scitranslmed.aaa3787

Bezerra, B.B., Silva, G.P.D.D., Coelho, S.V.A., Correa, I.A., Souza, M.R.M.D., Macedo, K.V.G., Matos, B.M., Tanuri, A., Matassoli, F.L., Costa, L.J.D., Hildreth, J.E.K., Arruda, L.B.D., 2022. Hydroxypropyl-beta-cyclodextrin (HP-BCD) inhibits SARS-CoV-2 replication and virus-induced inflammatory cytokines. Antiviral Res 205, 105373. 10.1016/j.antiviral.2022.105373

Biering, S.B., Akey, D.L., Wong, M.P., Brown, W.C., Lo, N.T.N., Puerta-Guardo, H., 2021. Structural basis for antibody inhibition of flavivirus NS1-triggered endothelial dysfunction. Science 371(6525):1, 194–200. 10.1126/science.abc0476.

Biering, S.B., Gomes De Sousa, F.T., Tjang, L.V., Pahmeier, F., Zhu, C., Ruan, R., Blanc, S.F., Patel, T.S., Worthington, C.M., Glasner, D.R., Castillo-Rojas, B., Servellita, V., Lo, N.T.N., Wong, M.P., Warnes, C.M., Sandoval, D.R., Clausen, T.M., Santos, Y.A., Fox, D.M., Ortega, V., Näär, A.M., Baric, R.S., Stanley, S.A., Aguilar, H.C., Esko, J.D., Chiu, C.Y., Pak, J.E., Beatty, P.R., Harris, E., 2022. SARS-CoV-2 Spike triggers barrier dysfunction and vascular leak via integrins and TGF-β signaling. Nat Commun 13, 7630. 10.1038/s41467-022-34910-5

Braga, S., Barbosa, J., Santos, N., El-Saleh, F., Paz, F., 2021. Cyclodextrins in Antiviral Therapeutics and Vaccines. Pharmaceutics 13, 409. 10.3390/pharmaceutics13030409

Coelho, D.R., Carneiro, P.H., Mendes-Monteiro, L., Conde, J.N., Andrade, I., Cao, T., Allonso, D., White-Dibiasio, M., Kuhn, R.J., Mohana-Borges, R., 2021. ApoA1 Neutralizes Proinflammatory Effects of Dengue Virus NS1 Protein and Modulates Viral Immune Evasion. J Virol 95, e01974–20. 10.1128/JVI.01974-20

Di Cagno, M., 2016. The Potential of Cyclodextrins as Novel Active Pharmaceutical Ingredients: A Short Overview. Molecules 22, 1. 10.3390/molecules22010001

Garrido, P.F., Calvelo, M., Blanco-González, A., Veleiro, U., Suárez, F., Conde, D., Cabezón, A., Piñeiro, Á., Garcia-Fandino, R., 2020. The Lord of the NanoRings: Cyclodextrins and the battle against SARS-CoV-2. Int J Pharm 588, 119689. 10.1016/j.ijpharm.2020.119689

Glasner, D.R., Puerta-Guardo, H., Beatty, P.R., Harris, E., 2018. The Good, the Bad, and the Shocking: The Multiple Roles of Dengue Virus Nonstructural Protein 1 in Protection and Pathogenesis. Annu Rev Virol 5, 227–253. 10.1146/annurev-virology-101416-041848

Hastings, C., Liu, B., Hurst, B., Cox, G.F., Hrynkow, S., 2022. Intravenous 2-hydroxypropyl-β-cyclodextrin (Trappsol® CycloTM) demonstrates biological activity and impacts cholesterol metabolism in the central nervous system and peripheral tissues in adult subjects with Niemann-Pick Disease Type C1: Results of a phase 1 trial. MolGenet Metab 137, 309–319. 10.1016/j.ymgme.2022.10.004

Hussein, H.A.M., Thabet, A.A., Wardany, A.A., El-Adly, A.M., Ali, M., Hassan, M.E.A., Abdeldayem, M.A.B., Mohamed, A.-R.M.A., Sobhy, A., El-Mokhtar, M.A., Afifi, M.M., Fathy, S.M., Sultan, S., 2024. SARS-CoV-2 outbreak: role of viral proteins and genomic diversity in virus infection and COVID-19 progression. Virol J 21, 75. 10.1186/s12985-024-02342-w

Jambhekar, S.S., Breen, P., 2016. Cyclodextrins in pharmaceutical formulations I: structure and physicochemical properties, formation of complexes, and types of complex. Drug Discov Today 21, 356–362. 10.1016/j.drudis.2015.11.017

Jones, S.T., Cagno, V., Janeček, M., Ortiz, D., Gasilova, N., Piret, J., Gasbarri, M., Constant, D.A., Han, Y., Vuković, L., Král, P., Kaiser, L., Huang, S., Constant, S., Kirkegaard, K., Boivin, G., Stellacci, F., Tapparel, C., 2020. Modified cyclodextrins as broad-spectrum antivirals. Sci Adv 6, eaax9318. 10.1126/sciadv.aax9318

Katzelnick, L.C., Coloma, J., Harris, E., 2017. Dengue: knowledge gaps, unmet needs, and research priorities. Lancet Infect Dis 17, e88–e100. 10.1016/S1473-3099(16)30473-X

Kovacs, T., Nagy, P., Panyi, G., Szente, L., Varga, Z., Zakany, F., 2022. Cyclodextrins: Only Pharmaceutical Excipients or Full-Fledged Drug Candidates? Pharmaceutics 14(12), 2559. 10.3390/pharmaceutics14122559

Li, Xinyu, Yuan, H., Li, Xiaozhen, Wang, H., 2023. Spike protein mediated membrane fusion during SARS-CoV-2 infection. J Med Virol 95, e28212. 10.1002/jmv.28212

Luo, Y., Zhang, Z., Ren, J., Dou, C., Wen, J., Yang, Y., Li, X., Yan, Z., Han, Y., 2024. SARS-Cov-2 spike induces intestinal barrier dysfunction through the interaction between CEACAM5 and Galectin-9. Front Immunol 15, 1303356. 10.3389/fimmu.2024.1303356

Matassoli, F.L., Leão, I.C., Bezerra, B.B., Pollard, R.B., Lütjohann, D., Hildreth, J.E.K., Arruda, L.B.D., 2018. Hydroxypropyl-Beta-Cyclodextrin Reduces Inflammatory Signaling from Monocytes: Possible Implications for Suppression of HIV Chronic Immune Activation. mSphere 3, e00497–18. 10.1128/mSphere.00497-18

Modhiran, N., Song, H., Liu, L., Bletchly, C., Brillault, L., Amarilla, A.A., Xu, X., Qi, J., Chai, Y., Cheung, S.T.M., Traves, R., Setoh, Y.X., Bibby, S., Scott, C.A.P., Freney, M.E., Newton, N.D., Khromykh, A.A., Chappell, K.J., Muller, D.A., Stacey, K.J., Landsberg, M.J., Shi, Y., Gao, G.F., Young, P.R., Watterson, D., 2021. A broadly protective antibody that targets the flavivirus NS1 protein. Science 371, 190–194. 10.1126/science.abb9425

Modhiran, N., Watterson, D., Blumenthal, A., Baxter, A.G., Young, P.R., Stacey, K.J., 2017. Dengue virus NS1 protein activates immune cells via TLR4 but not TLR2 or TLR6. Immunol Cell Biol 95, 491–495. 10.1038/icb.2017.5

Modhiran, N., Watterson, D., Muller, D.A., Panetta, A.K., Sester, D.P., Liu, L., Hume, D.A., Stacey, K.J., Young, P.R., 2015. Dengue virus NS1 protein activates cells via Toll-like receptor 4 and disrupts endothelial cell monolayer integrity. Sci Transl Med 7, 304ra142. 10.1126/scitranslmed.aaa3863

Orozco, S., Schmid, M.A., Parameswaran, P., Lachica, R., Henn, M.R., Beatty, R., Harris, E., 2012. Characterization of a model of lethal dengue virus 2 infection in C57BL/6 mice deficient in the alpha/beta interferon receptor. J Gen Virol 93, 2152–2157. 10.1099/vir.0.045088-0

Pahmeier, F., Monticelli, S. R., Feng, X., Hjorth, C. K., Wang, A., Kuehne, A. I., Bakken, R. R., Batchelor, T. G., Lee, S. E., Middlecamp, M., Stuart, L., Duarte-Neto, A. N., Abelson, D. M., McLellan, J. S., Biering, S. B., Herbert, A. S., Chandran, K., & Harris, E., 2025. Antibodies targeting Crimean-Congo hemorrhagic fever virus GP38 limit vascular leak and viral spread. Sci Transl Med, 17(786), eadq5928. 10.1126/scitranslmed.adq5928

Panigrahi, S., Goswami, T., Ferrari, B., Antonelli, C.J., Bazdar, D.A., Gilmore, H., Freeman, M.L., Lederman, M.M., Sieg, S.F., 2021. SARS-CoV-2 Spike Protein Destabilizes Microvascular Homeostasis. Microbiol Spectrum 9, e00735–21. 10.1128/Spectrum.00735-21

Pardeshi, C.V., Kothawade, R.V., Markad, A.R., Pardeshi, S.R., Kulkarni, A.D., Chaudhari, P.J., Longhi, M.R., Dhas, N., Naik, J.B., Surana, S.J., García, M.C., 2023. Sulfobutylether-β-cyclodextrin: A functional biopolymer for drug delivery applications. Carbohydrate Polymers 301, 120347. 10.1016/j.carbpol.2022.120347

Puerta-Guardo, H., Biering, S. B., Castillo-Rojas, B., DiBiasio-White, M. J., Lo, N. T., Espinosa, D. A., Warnes, C. M., Wang, C., Cao, T., Glasner, D. R., Beatty, P. R., Kuhn, R. J., & Harris, E., 2024. Flavivirus NS1-triggered endothelial dysfunction promotes virus dissemination. Preprint, BioRxiv 2024; 2024.11.29.625931. 10.1101/2024.11.29.625931

Puerta-Guardo, H., Glasner, D.R., Espinosa, D.A., Biering, S.B., Patana, M., Ratnasiri, K., Wang, C., Beatty, P.R., Harris, E., 2019. Flavivirus NS1 Triggers Tissue-Specific Vascular Endothelial Dysfunction Reflecting Disease Tropism. Cell Rep 26, 1598–1613.e8. 10.1016/j.celrep.2019.01.036

Puerta-Guardo, H., Glasner, D.R., Harris, E., 2016. Dengue Virus NS1 Disrupts the Endothelial Glycocalyx, Leading to Hyperpermeability. PLoS Pathog 12, e1005738. 10.1371/journal.ppat.1005738

Raïch-Regué, D., Tenorio, R., Fernández de Castro, I., Tarrés-Freixas, F., Sachse, M., Perez-Zsolt, D., Muñoz-Basagoiti, J., Fernández-Sánchez, S.Y., Gallemí, M., Ortega-González, P., Fernández-Oliva, A., Gabaldón, J.A., Nuñez-Delicado, E., Casas, J., Roca, N., Cantero, G., Pérez, M., Usai, C., Lorca-Oró, C., Alert, J.-V., Segalés, J., Carrillo, J., Blanco, J., Clotet Sala, B., Cerón-Carrasco, J.P., Izquierdo-Useros, N., Risco, C., 2023. β-Cyclodextrins as affordable antivirals to treat coronavirus infection. Biomed Pharmacother 164, 114997. 10.1016/j.biopha.2023.114997

Rathore, A.P.S., Paradkar, P.N., Watanabe, S., Tan, K.H., Sung, C., Connolly, J.E., Low, J., Ooi, E.E., Vasudevan, S.G., 2011. Celgosivir treatment misfolds dengue virus NS1 protein, induces cellular pro-survival genes and protects against lethal challenge mouse model. Antiviral Res 92, 453–460. 10.1016/j.antiviral.2011.10.002

Scaturro, P., Cortese, M., Chatel-Chaix, L., Fischl, W., Bartenschlager, R., 2015. Dengue Virus Non-structural Protein 1 Modulates Infectious Particle Production via Interaction with the Structural Proteins. PLoS Pathog 11, e1005277. 10.1371/journal.ppat.1005277

Senti, G., Iannaccone, R., Graf, N., Felder, M., Tay, F., Kündig, T., 2013. A randomized, double-blind, placebo-controlled study to test the efficacy of topical 2-hydroxypropyl-Beta-cyclodextrin in the prophylaxis of recurrent herpes labialis. Dermatol 226, 247–52. 10.1159/000349991

Sousa, F.T.G.D., Biering, S.B., Patel, T.S., Blanc, S.F., Camelini, C.M., Venzke, D., Nunes, R.J., Romano, C.M., Beatty, P.R., Sabino, E.C., Harris, E., 2022. Sulfated β-glucan from Agaricus subrufescens inhibits flavivirus infection and nonstructural protein 1-mediated pathogenesis. Antiviral Res 203, 105330. 10.1016/j.antiviral.2022.105330

Watanabe, S., Rathore, A.P.S., Sung, C., Lu, F., Khoo, Y.M., Connolly, J., Low, J., Ooi, E.E., Lee, H.S., Vasudevan, S.G., 2012. Dose- and schedule-dependent protective efficacy of celgosivir in a lethal mouse model for dengue virus infection informs dosing regimen for a proof of concept clinical trial. Antiviral Res 96, 32–35. 10.1016/j.antiviral.2012.07.008

Yu, S., Hu, H., Ai, Q., Bai, R., Ma, K., Zhou, M., Wang, S., 2023. SARS-CoV-2 Spike-Mediated Entry and Its Regulation by Host Innate Immunity. Viruses 15(3), 639. 10.3390/v15030639

